# The application of text mining algorithms in summarizing trends in anti-epileptic drug research

**DOI:** 10.1101/269308

**Authors:** Shatrunjai P. Singh, Swagata Karkare, Sudhir M. Baswan, Vijendra P. Singh

**Author notes:** Corresponding Author: Shatrunjai P. Singh.

## Abstract

Content summarization is an important area of research in traditional data mining. The volume of studies published on anti-epileptic drugs (AED) has increased exponentially over the last two decades, making it an important area for the application of text mining based summarization algorithms. In the current study, we use text analytics algorithms to mine and summarize 10,000 PubMed abstracts related to anti-epileptic drugs published within the last 10 years. A Text Frequency – Inverse Document Frequency based filtering was applied to identify drugs with highest frequency of mentions within these abstracts. The US Food and Drug database was scrapped and linked to the results to quantify the most frequently mentioned modes of action and elucidate the pharmaceutical entities marketing these drugs. A sentiment analysis model was created to score the abstracts for sentiment positivity or negativity. Finally, a modified Latent Dirichlet Allocation topic model was generated to extract key topics associated with the most frequently mentioned AEDs. Results of this study provide accurate and data intensive insights on the progress of anti-epileptic drug research.

## 2. Introduction

The unparalleled surge in published biomedical literature has made it difficult to define quantitative and qualitative summarization of a specific topic. Recent advances in computational power have led to an increase in the use of text mining approaches to facilitate the summarization and content review (Khordad and Mercer, 2017; Moradi and Ghadiri, 2017; Zhu et al., 2013). Open source analytical tools can rapidly ingest vast sources and volumes of information which can then be further pipelined into key insights using algorithms like feature extraction, topic modeling and sentiment analysis, allowing accurate summarization (Mishra et al., 2014). These text mining approaches have already been employed in analyzing a wide array of topics like oncology databases (Zhu et al., 2013), impact of financial crises on suicides (Jung et al., 2017), awareness of climate change in rural communities (Bell, 2013) and analyzing the sentiment of diabetes patients on the twitter platform (Salas-Zárate et al., 2017). Although text mining has been employed in several domains of biomedical research, its use remains infrequent in many important therapeutic areas, including neuroscience research (Singh, 2015).

Epilepsy, the fourth most frequent neurological disorder, affects more than sixty million people globally (Singh, 2015; Singh and Karkare, 2017; Trinka et al., 2015; Singh et al., 2013, 2015). The social stigma linked with this condition often primes depression and is frequently associated with a decline in the quality of life (Benson et al., 2016; Hester et al., 2016; Luna et al., 2017).

The problem is exacerbated by the confusion of focusing research efforts on multiple anti-epileptic drugs (AEDs), some of which show mixed results in the refractory epileptic populations (Ahmad et al., 2017; de Biase et al., 2017; Nolan et al., 2015; Pellock et al., 2017; Turner and Perry, 2017). The sheer volume of new research on AEDs cripples any meaningful insight generation.

In this study, we analyze 10,000 PubMed abstracts related to AEDs with the end goal of content summarization and insight generation. Abstracts containing US FDA curated list of drugs were identified and analyzed for drug frequency, mode of action and the pharmaceutical entities manufacturing the most frequent drugs were acknowledged. A modified latent dirichlet analysis algorithm with a bigram tokenizer (mLDA) was used to extract key topics discussed in these abstracts. Finally, sentiment analysis was utilized to analyze which of these anti-epileptic drugs are promising candidates for further research based on associated positive sentiments.

## 3. Methods

### 3.1 Data Collection

We used an R-software based PubMed scrapper to download 10,000 abstracts positive either of these keywords: ‘anti-epileptic drugs’, ‘anti-convulsant drugs’ and ‘AED’. Only abstracts published between 01/01/2007 to 01/01/2017 were included in the study. Papers with no abstracts or written in languages other than English were filtered out. The raw abstract data was uploaded to a public repository for open access (Singh and Karkare, 2018). The raw R code used for the analysis was deposited in a *Githib* repository (https://github.com/shatrunjai/aed_pubmed). A document term matrix (DTM) was created from the abstracts and was compared to the list of drugs approved in the last decade, obtained from the US Food and Drug Administration website (https://www.fda.gov/Drugs.htm). Only abstracts focusing on at least one of these drugs was included for further analysis.

### 3.2 Data Processing

Collected abstracts were scrubbed for numbers, non-English characters and stop words. The Stanford stop words list was used as the default stop word repository (https://nlp.stanford.edu/IR-book/html/htmledition/dropping-common-terms-stop-words-1.html). Stemming of abstract was conducted according to the Porter stemmer (Porter, 1980). A document term matrix was created as described in Stanford NLP (https://nlp.stanford.edu/).A Term-Frequency-Inverse Document Frequency (TF-IDF) matrix was created and further frequency calculations were performed only on relevant TF-IDF terms as described in (Jones, 1972).The frequency matrix had a mean word frequency of 272 words and a standard deviation of 17 words. Words with frequency cut off two standard deviations from the mean word frequency were filtered from the list.

### 3.3 Modified LDA based Topic Modelling

Latent Dirichlet Allocation (LDA) is a well-defined, unsupervised, generative, probabilistic method for modeling data and is frequently used in topic modeling (Blei et al., 2003). We created a modified Latent Dirichlet Allocation (mLDA) algorithm which assumes that each document can be denoted as a probabilistic distribution over latent topics and that the topic distribution in all documents share a common Dirichlet prior distribution. We also included a bigram tokenizer to better represent scientific abstracts. Each latent topic in the mLDA model is also represented as a probabilistic model over words and the word distributions of topics share a common Dirichlet prior distribution as well. Given a corpus *M* consisting of *N* documents, with document *d* having *K*_*d*_ words (*d* **∈**{1,…, *N*}), mLDA models *M* according to the following generative process (Blei et al., 2003; Li et al., 2016):

a. Select a multinomial distribution *φ*_*t*_ for topic *t* (*t* **∈**{1,…, *T*}) from a Dirichlet prior distribution with parameter *β*.
b. Select a multinomial distribution *θ*_*d*_ for document *d*(*d* **∈**{1,…, *N*}) from a Dirichlet prior distribution with parameter *α*.
c. For a word *w*_*n*_(*n* **∈**{1,…, *K*_*d*_}) in document *d*,

i. Select a topic *Z*_*n*_ from *θ*_*d*_.
ii. Select a word *W*_*n*_ from *φ*_*zn*_.

This generative process has words in documents are the only detected variables whereas others are latent variables (*φ* and *θ*) and hyper parameters (*α* and *β*). In order to deduce the latent variables and hyper parameters, the probability of experiential data *M* is calculated as follows:

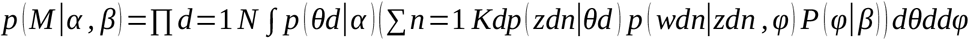

Due to the coupling between *θ* and *φ* in the integrand (above equation), the precise implication in mLDA is obstinate (Blei et al., 2003).The number of topics was selected according to the Rate of Perplexity Change (RPC) previously described by Zhao and colleagues (Zhao et al., 2015). This algorithm yielded two key topics on average which were further curated manually.

### 3.4 Sentiment Analysis

To evaluate sentiment for each abstract, the *Sentiment Analysis* and *Tm* libraries were used within R-software, Version 0.98.109 ([CSL STYLE ERROR: reference with no printed form.]). *Sentiment Analysis* algorithm is a well-established sentiment analysis (SA) protocol and has been cited by over a 1000 journal publications according to google scholar. *Sentiment Analysis* and *Tm* packages assign three sentiment scores (“positive,” “negative,” and “neutral”) to each word, based on a generalized classification system developed by the authors which uses a combination of human-annotated and Artificial Intelligence based sentiment scoring algorithms (Bagheri and Islam, 2017). Further, we employed the “bag-of-words” approach which has been established to be very dependable for document-level SA, with aggregate-level performance approximately equivalent to more refined methods (Gayle and Shimaoka, 2017).

For the current study, nouns were excluded from the analysis as they contain little to no information (Pinheiro et al., 2015). The sentiment of each abstract was calculated by combining the scores of all pertinent word tokens. A sentiment score ranging from −1 to +1 was allocated for each abstract based on the assessed grade of negative or positive sentiment. For further analysis and visualization, unstandardized scores were normalized to a distribution with a mean of zero (x^−^=0) and standard deviation of one (*σ*_x^−^_=1). All abstracts were assigned values of ‘positive’ (score>+1),‘negative’ (score>−1) or ‘neutral’ (−1<score<+1).

### 3.5 Machine Learning

The *Sentiment Analysis* package uses a one class support vector machine (SVM) algorithm to classify the expressions and phrases within the abstracts based on Stanford core NLP trained algorithm. SVM is a supervised analytical method that classifies based on the degree to which the several input cases (i.e., expression vectors) predict a given binary class, like the presence of absence of positive sentiment (Salas-Zárate et al., 2017). All input terms, i.e. the bigrams can thus be assessed in terms of “importance” with respect to a given label (Gayle and Shimaoka, 2017).The classifier was retrained on a 7000-abstract sample curated dataset optimized for misclassification rate, precision and recall metrics.

### 3.6 Statistical Analyses

Microsoft SQL Server (version 2012) was used to query the dataset for different clone compositions, and statistical analysis was performed using R-statistical software (Version 0.98.109). Significance was determined using a two-tailed Student’s t-test for data that met assumptions of normality and equal variance. The Mann-Whitney rank sum test was used for non-normal data. Proportions were compared using z-tests. Values presented are means ± SEM or medians [range], as appropriate. The experiment-wise error was conservatively set at 0.001 (Cumming, 2010). Corrections for multiple comparisons were done using a Bonferroni correction.

### 3.7 Figure Preparation

The results from R-software were exported into csv files which were imported into Tableau (version 8.0) or Microsoft Excel (version 2013) which were then used to create graphs and visualizations. Tables were created in Microsoft Word (version 2013).

## 4. Results

### 4.1 Characterizing the most published anti-epileptic drugs in the last 10 years

To study AEDs that appeared in PubMed abstracts (2007-2017), an R scrapper was used to parse 10,000 PubMed abstracts. To identify abstracts specifically related to AEDs, this scrapped dataset was cross-referenced with the United States Food and National Drug database (US FDA) of drugs. A total of 130 drugs (Figure 1) with a mean of 69.34 abstracts per drugs and a standard deviation of 22.03 abstracts per drugs were identified. The top 5 most frequent drugs were: Gabapentin (abstract count=1371, Figure 1), Levetiracetam (abstract count=1304, Figure 1), Topiramate (abstract count=1027, Figure 1), Lamotrigine (abstract count=989, Figure 1) and Acetazolamide (abstract count=518, Figure 1). A year-by-year frequency of selected drug abstracts was performed for all the drugs beginning the year 1980 (Figure 2) to follow their research trends.

**Figure 1.**
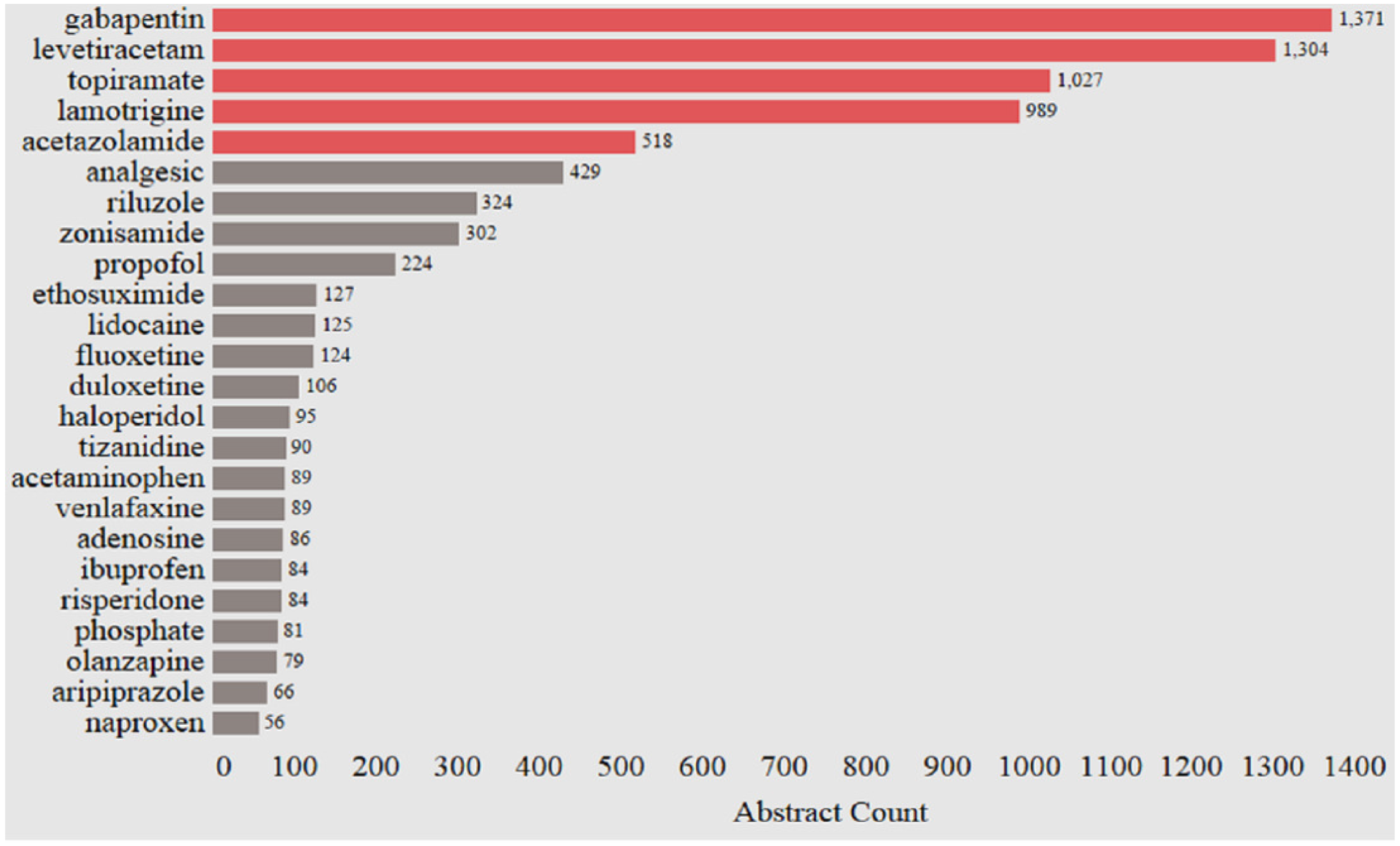
The frequency of PubMed abstracts with anti-epileptic drug mentions from 2007 to 2017. Shown here are the drugs with more than 50 abstracts. The AED’s further analyzed are in red.

**Figure 2.**
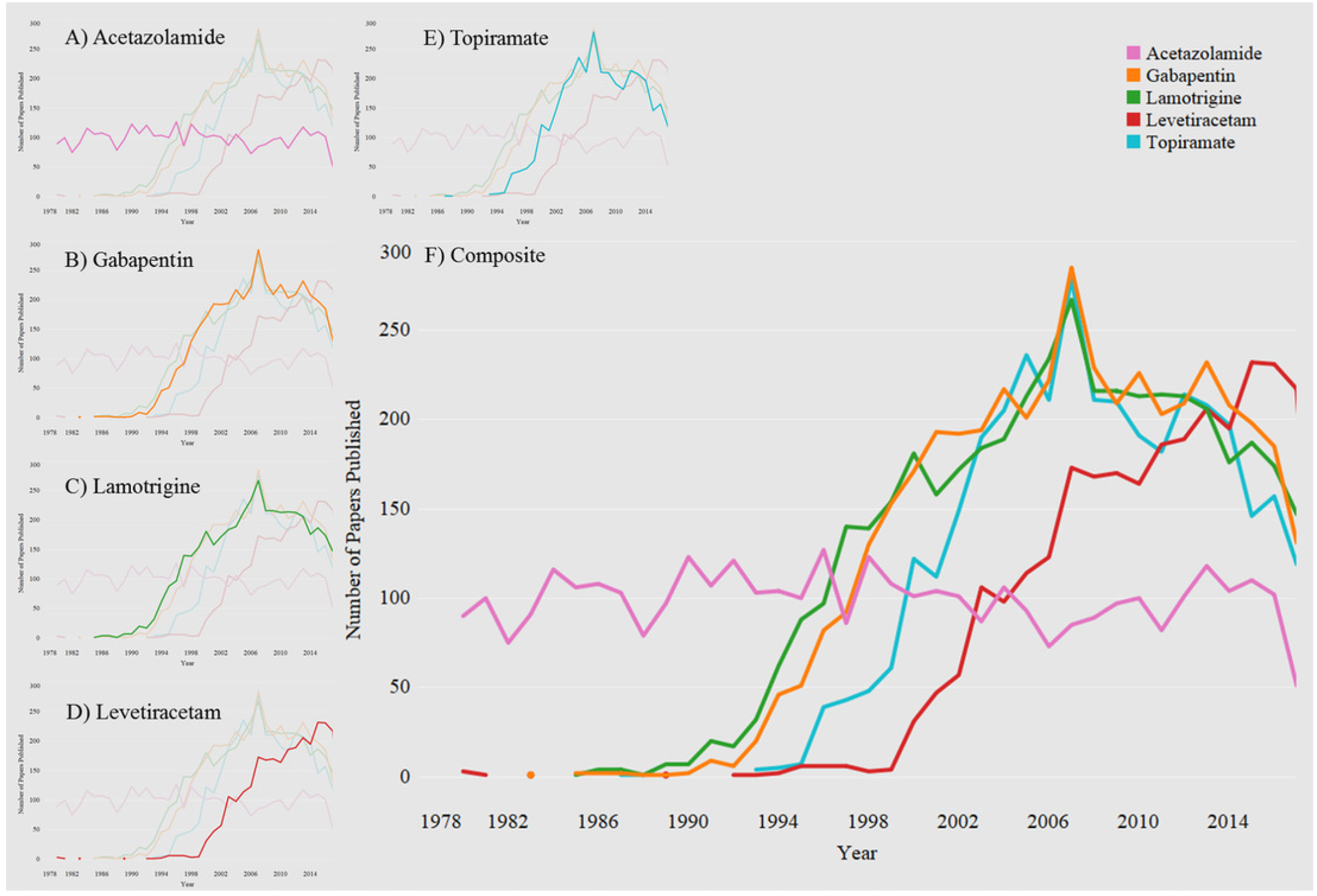
Yearly mentions of the five most frequently occurring anti-epileptic drugs. The frequency of abstracts published between 1980 to 2017 for A) Acetazolamide, B) Gabapentin, C) Lamotrigine, D) Levetiracetam and E) Topiramate and F) Composite of all the drugs (A through E).

### 4.2 Characterizing Drug Class of the most published drugs

For all drugs, their pharmaceutical drug categorization was evaluated by using FDA definitions (https://www.fda.gov/drugs/informationondrugs/ucm079436.htm). As expected, Anti-Epileptic agents and CNS activity suppression agents were at the top of the list of our drug matches (Figure 3). However, cox-2 inhibitors, mood stabilizers, cytochrome p450-2C19 inhibitors, analgesics and serotonin reuptake inhibitors also frequent in the class of researched AEDs (Figures 3), reflecting the diversity in research initiatives.

**Figure 3.**
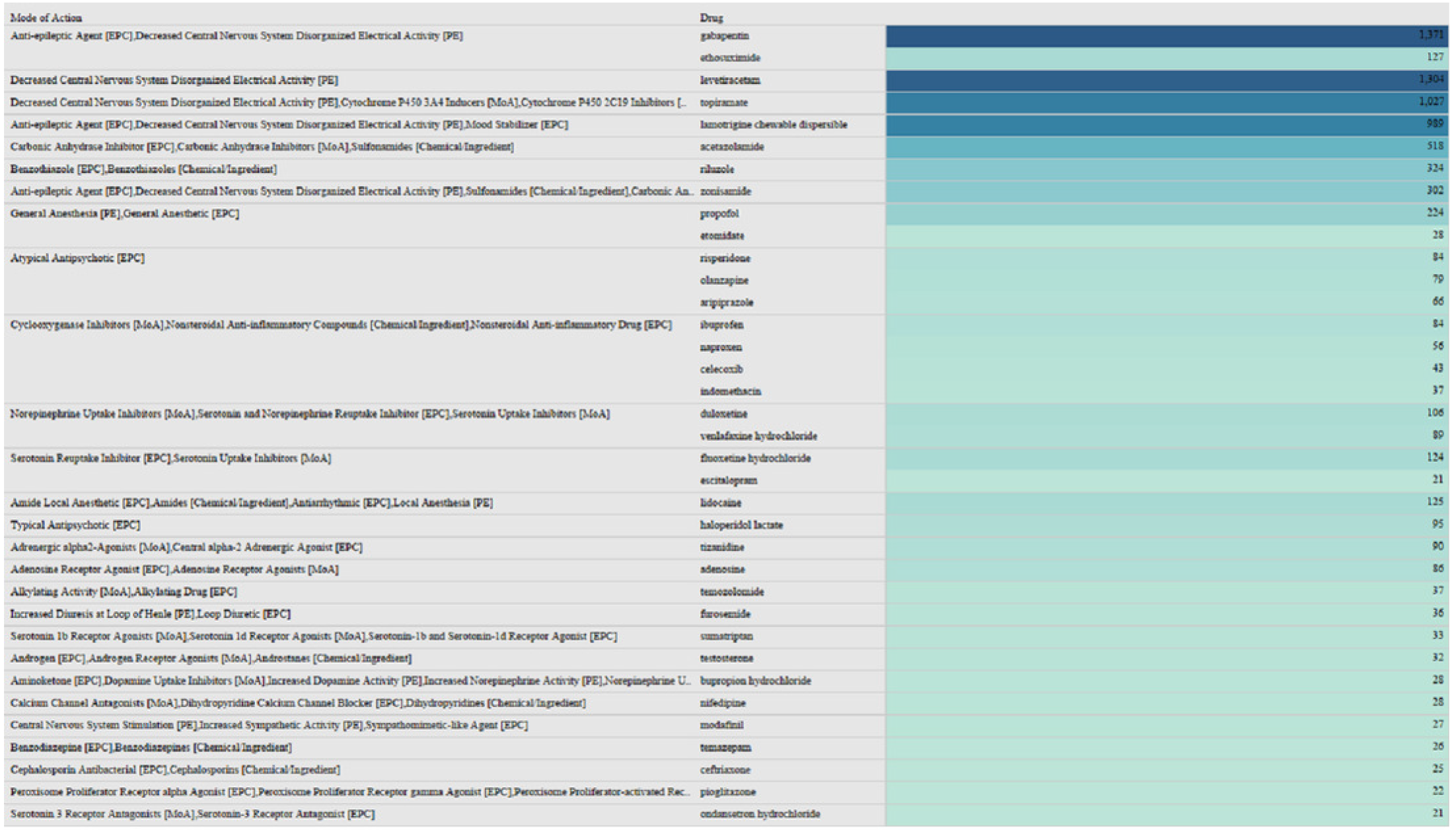
The most common drug classes found in the abstract with the frequency of their occurrence. Drug classification was obtained from the US FDA classification.

### 4.3 Characterizing Pharmaceutical Industries with the most published drugs

Next, the pharmaceutical companies associated with the highest frequency of drug mentions in the 10,000 abstracts selected for the study were extracted (Figure 4). Some companies had more than 5 drugs (Sagent Pharmaceuticals and Zydus Pharmaceuticals Inc. with 14 and 9 drugs, respectively). Zydus Pharmaceutical’s Topiramate along with the other 8 drugs appears to lead the list in terms of the number of drugs and the frequency of abstract mentions. However, other companies like A-S Medication Solutions which despite having only one drug (Gabapentin), were still top-ranked in abstract mention frequency (Figure 4).

**Figure 4.**
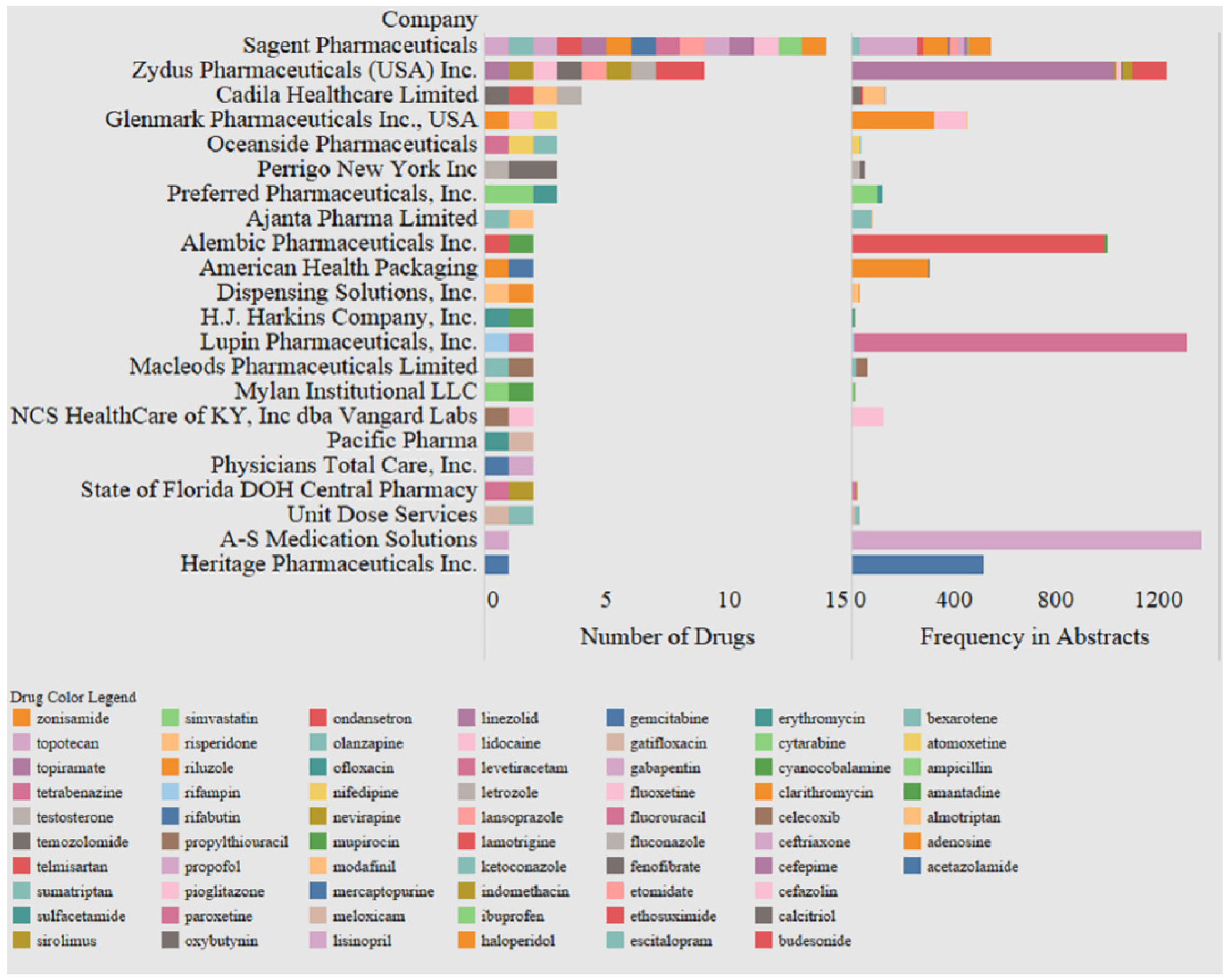
Pharmaceutical companies which have the most number of drugs mentioned in the last 10000 abstracts. The registered name of the company is shown in the first pane, the number of drugs that made it into the study criterion in the second pane and the frequency of drug appearance in the abstracts in the third pane.

### 4.4 Using sentiment analysis to score the abstracts with the top anti-epileptic drugs

A sentiment analysis was performed on all the abstracts containing the keyword ‘anti-epileptic drugs’ or ‘AED’ or ‘anti-convulsion drugs’. An initial analysis revealed a strong correlation between negative sentiment and the frequency of abstract mention (Table 1, correlation coefficient=0.68). To correct for this, a normalized sentiment score (Sentiment-Sentiment _mean_/Sentiment _s.d._) was calculated for each drug (Table 1). The sentiment value/abstract correlation was manually tested for accuracy. Drugs Lisinopril (normalized sentiment score= −3.0) and Telmisartan (normalized sentiment score= 3.0) had the highest positive normalized sentiment of all the drugs, indicating that these appeared in abstracts with positive connotations (‘positive outcome’, ‘no side effects’) more often than other drugs. Conversely, Ethosuximide (normalized sentiment score= −0.9) and Meloxicam (normalized sentiment score= −2.3) had the most negative sentiment, indicating appearance in abstracts with negative connotations (‘negative outcomes’, ‘side effects’).

**Table 1.**
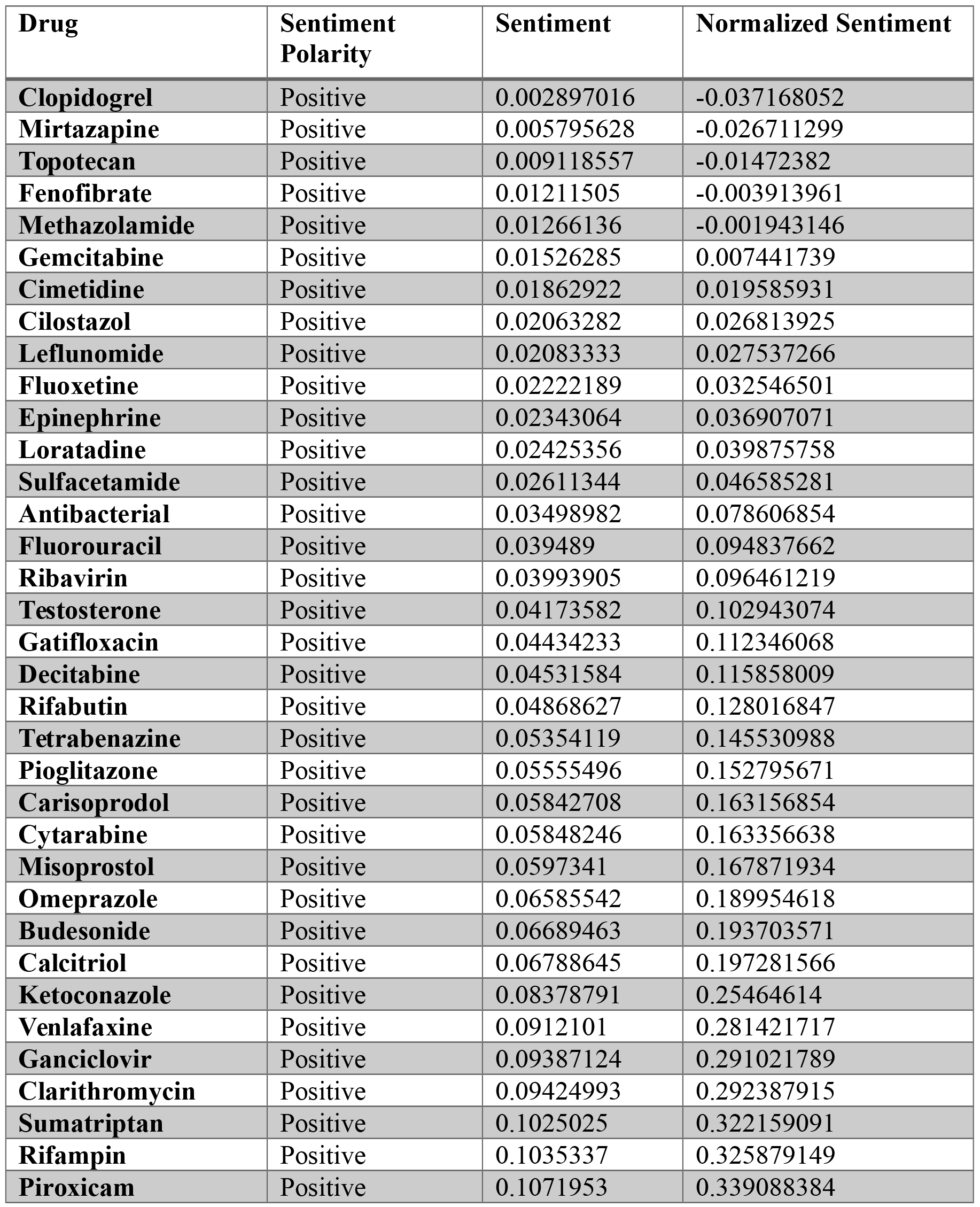
Sentiment analysis on abstracts with AED mentions. The sentiment of all the abstracts (not-normalized) is shown as either positive (score>0, green) or negative (score<0, red). The normalized sentiment scores (score-mean/S.D.) is shown in the right most column. The mean sentiment score was 0.01320 and the standard deviation was 0.2772. ?Md

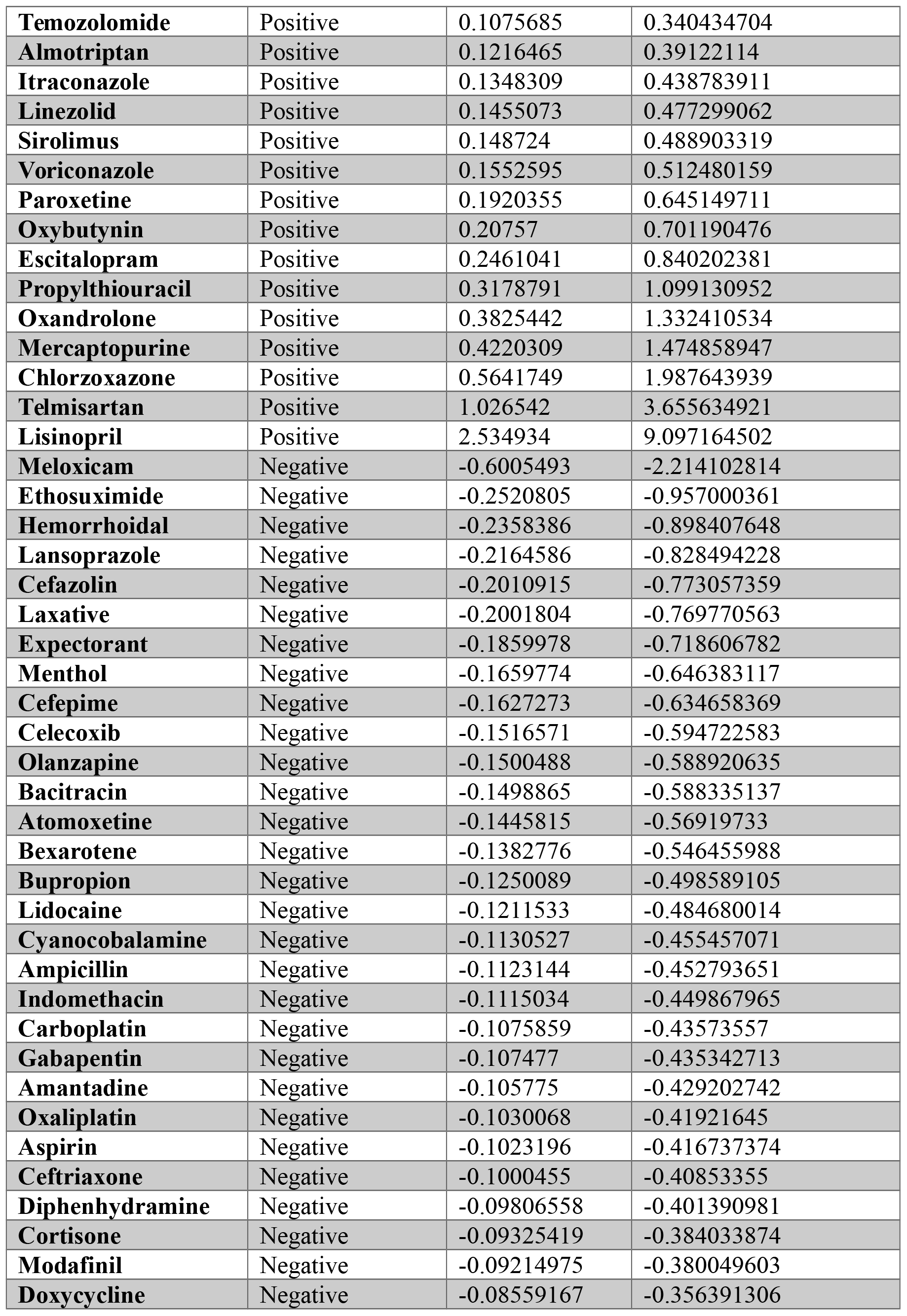

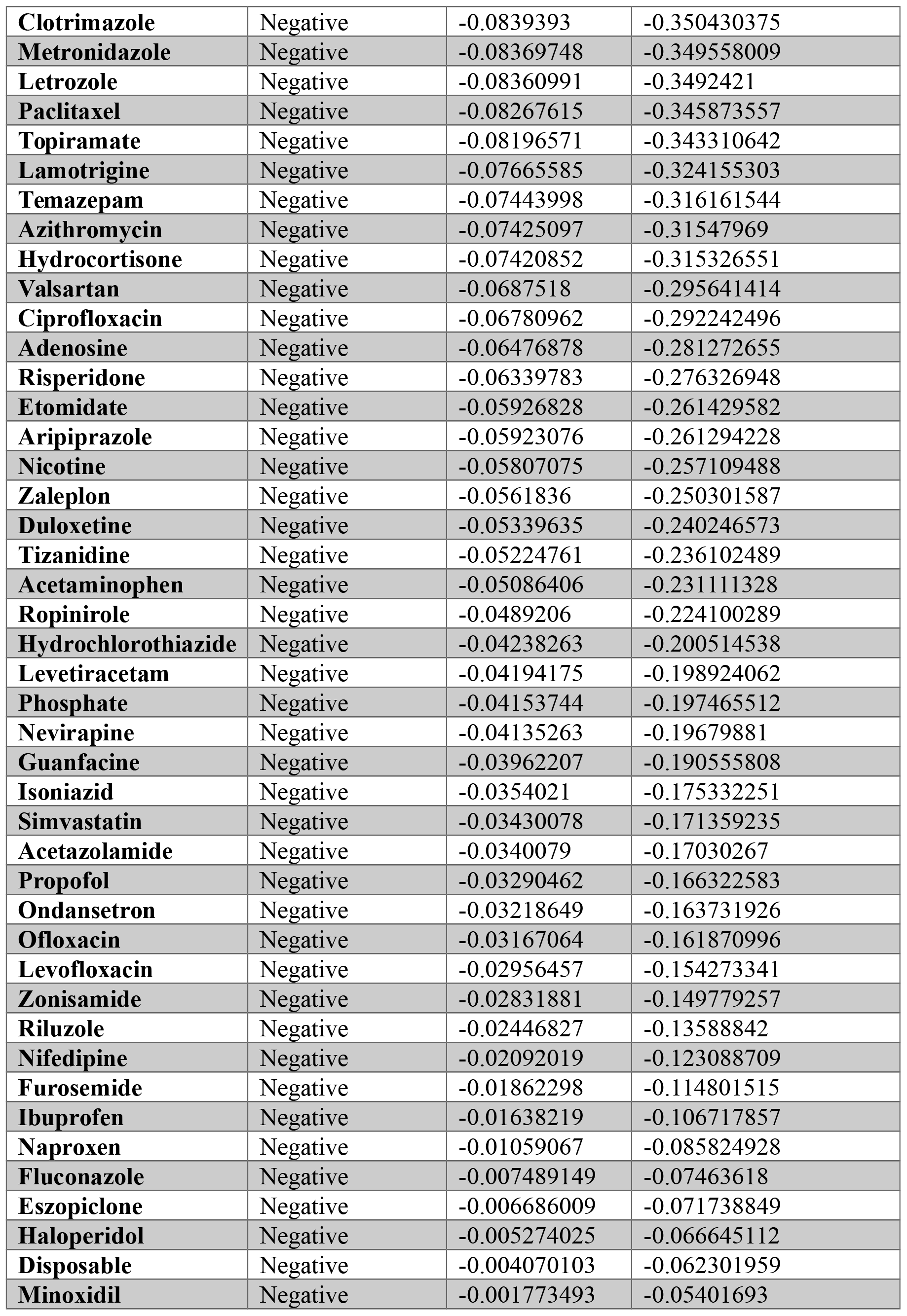

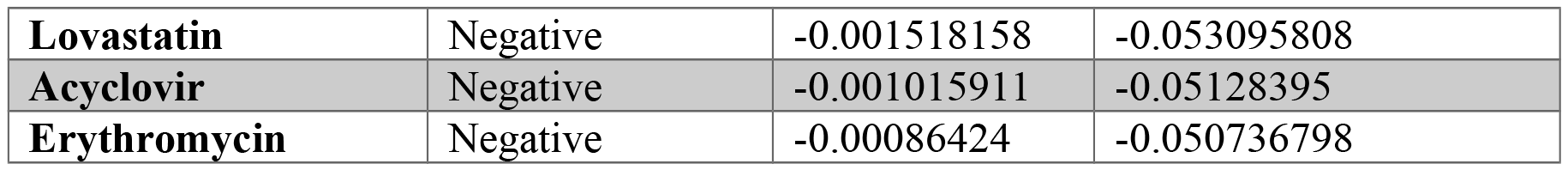

### 4.5 A modified Latent Dirichlet Algorithm reveals topics associated with the top 5 most mentioned anti-epileptic drugs

An mLDA algorithm was employed to identify the key topics being discussed in the papers associated with the top 5 drug mentions (Table 2). Key words associated with the top topic indicated research on the lines of spinal surgery and pain outcomes. Levetiracetam was associated with topics including its use in refractory and generalized seizure, response bias by gender and its association with Brivaracetam. Topiramate was associated with topics including long term side effects, the development of drug-resistance, and its effect on Lennox-Gastaut syndrome. Acetazolamide was associated with one topic indicating research on its effect on visual acuity and macular degeneration. Finally, Lamotrigine was associated with one topic indicating possible side effects of dry mouth and blood spots at higher concentration of the drug.

**Table 2.** Results from mLDA based Topic models run on abstract containing the top 5 drugs. Each topic is represented by the top keywords defining the topic.

| Gabapentin |  | Levetiracetam |  | Topiramate |  | Acetazolamid<br>e | Lamotrigine |
| --- | --- | --- | --- | --- | --- | --- | --- |
| Topic 1 | Topic 2 | Topic 1 | Topic 2 | Topic 1 | Topic 2 | Topic 1 | Topic 1 |
| Pain | Analog | Refractory | Indicated | LGS | Controllin<br>g | Clinical | Blood |
| Group | Incidence | Generalized | Woman | Epilepsy | Drug<br>Resistant | Macular | Dried |
| Controlled | Scores | Seizures | Partial<br>Epilepsy | Lennoxgastau<br>t | Effective | Visual | Spots |
| Spinal | Inclusion | New | Response | Drugs | Long Term | Acuity | Concentrations |
| Surgery | Scale | Brivaracetam | Therapy | Resolution | Drop | Better | Accuracy |

## 5. Discussion

In this study, we use text analytics algorithms to summarize the latest development in anti-epileptic drug research. We mined the top five drugs that have been extensively published in PubMed, elucidate the pharmaceutical entities manufacturing/marketing these drugs, and also provided sentiment based direction on how this research is trending. Finally, we created an mLDA based topic modelling algorithm to discuss key topics associated with these drugs.

The most popular AED’s conventionally used as first line treatment include primidone, ethosuximide, benzodiazepines, carbamazepine and phenobarbital. In the last 20 years, the Food and Drug Administration (FDA) has further approved twelve new AED’s and have a longer list of these drugs in the clinical trial pipelines (Asconapé, 2010). Although all of these compounds have been used to treat epilepsy for more than a century a true anti-epileptic drug effective against all seizure grades and all demographics is still unavailable and approximately 30% of patients with epilepsy do not respond to any existing AEDs (Glauser et al., 2006; Singh, 2015). This has fueled basic research into new pharmacological agents with better safety and tolerability, ease of use and better titration rate, fewer potential interactions, and increased efficacy in comorbidities (Azar and Abou-Khalil, 2008). The resultant research from studies on different aspects of multiple AEDs has often made research summarization difficult and calls for newer computational approaches.

PubMed, the most extensively used warehouse of biomedical literature comprises of more than 20 million abstracts and is increasing at a frequency of over 90,000 abstracts per year: the quantity of articles added each year to PubMed has increased three times in the last 10 years (Andronis et al., 2011). As research on a solitary subject may extend across numerous scientific areas and technical journals, it is progressively problematic for scientists to trail all advances in their area of work. The dispersal of information to many different journals and scientific subgroups has created and ‘islets of scientific knowledge’ and has led to the improvement of literature mining approaches pointing to link ideas and opinions that are not cited in the same editorial. The process of deducing implied knowledge from apparently unrelated concepts has been named literature-based discovery (LBD) (Andronis et al., 2011). These LBD methods have been used in the past for the purpose of theory ideation in association with drug discovery. Some of these LBD techniques include PubMed text mining, TF-IDF based keyword generation, unsupervised document clustering, literature modelling, sentiment analysis and topic modelling techniques. In the current study, we use a subset of these techniques for AED centered research summarization.

The most frequently studied AED was found to be Gabapentin, which is indicated for the treatment of postoperative neuralgia in adults and for treating partial onset seizures in both pediatric and adult patients (Goa and Sorkin, 1993). Although the exact mode of Gabapentin action is unknown, it has been suggested that its activity depends on its interaction with voltage-gated calcium channels (Goa and Sorkin, 1993). Interestingly, topic modeling revealed the keywords ‘pain’ and ‘spinal surgery’ to be associated with this drug. However, although gabapentin is commonly used in pain management, its use in post-operative pain and spinal surgery is controversial (Chang et al., 2014; Yu et al., 2013).

Levetiracetam, the second most commonly researched anti-epileptic drug, is indicated as an adjunctive therapy in the treatment of partial onset seizures in patients ≥16 years of age with epilepsy(Deshpande and Delorenzo, 2014; Zheng et al., 2015). The precise mechanism(s) by which Levetiracetam exerts its antiepileptic effect is unknown, but studies suggest that this agent acts as a neuromodulator and treats seizures by inhibiting presynaptic calcium channels (Deshpande and Delorenzo, 2014). Topic modeling from this study revealed recent efforts towards comparing the efficacy of Levetiracetam to Brivaricetam which has been a topic of increasing interest over the year (Crepeau and Treiman, 2010; Lyseng-Williamson, 2011).

Topiramate is used as a monotherapy in children of ages two and above and as an adjunctive therapy for adults. Its use is children is specifically indicated for seizures related with Lennox-Gastaut syndrome (LGS) (Crumrine, 2011; Donegan et al., 2015; Hoy, 2016). Topic modeling showed a strong association of this agent with Lennox-Gastaut syndrome, a disorder which initiates seizures in children(Crumrine, 2011; VanStraten and Ng, 2012).

Acetazolamide, a carbonic anhydrase inhibitor is indicated for the treatment of centrencephalic epilepsies (petit mal, unlocalized seizures) and is also a popular drug for the treatment of glaucoma (Reiss and Oles, 1996; Millichap and Aymat, 1967). Results of the topic modeling used in this study support a strong association of this drug with keywords like ‘macular’, ‘visual’, ‘acuity’, all of which are glaucoma-related terms referring to the discovery of its anti-epileptic properties during treatment of glaucoma patients (Lyall, 2008). Lamotrigine is an antiepileptic drug indicated as an adjunctive therapy in children above the ages of two specifically for primary generalized tonic-clonic seizures. It is also indicated for the treatment of bipolar disorder in patients (Ramaratnam et al., 2016). Although the mechanism of action of this drug is unknown, in vitro pharmacological studies suggest that lamotrigine inhibits voltage-sensitive sodium channels, thereby stabilizing neuronal membranes and consequently modulating presynaptic transmitter release of excitatory amino acids (e.g., glutamate and aspartate). Topic modeling revealed the association of this drug with the terms ‘dried blood spots’, which suggests that research efforts have been focused on evaluating the safety profile of this drug, specifically in causing blood dyscrasias (Krasowski and McMillin, 2014; Milosheska et al., 2015; Baswan et al., 2016).

Sentiment analysis suggests that despite these drugs being well-established and approved lines of therapy in the treatment of a variety of epilepsies and seizures, all 5 drugs were associated with a negative sentiment. This indicates the possibility of mixed results in at least a subset of these research studies. Further, these findings suggest potential unmet need in the area of epilepsy treatment due to the dearth of positive sentiments surrounding these pharmacological agents.

## 6. Conclusion

This study demonstrates that although research efforts surrounding anti-epileptic treatments are moving in the right direction, there is an unmet need when it comes to the associated sentiments of researchers towards the most frequently studied agents. Despite the potential utility of these drugs in the treatment of epilepsy, their use in treatment could be hindered due to associated negative sentiments. Even though this study delineates the key topics surrounding AED research in the last decade, further research efforts should be conducted to understand the causal relationship between the negative sentiments and the pharmacological profile of these agents. Understanding these causative efforts can help lead the way for pharmaceutical manufacturers to devote research efforts towards improving the profiles of their drugs to better suit the needs of the patients.

